# Basic reproduction numbers of three strains of mouse hepatitis viruses in mice

**DOI:** 10.1101/2021.09.24.461643

**Authors:** Masataka Nakayama, Shigeru Kyuwa

## Abstract

Mouse hepatitis virus (MHV) is a murine coronavirus and one of the most important pathogens in laboratory mice. Although various strains of MHV have been isolated, they are generally excreted in the feces and transmitted oronasally via aerosols and contaminated bedding. In this study, we attempted to determine the basic reproduction numbers of three strains of MHV to improve our understanding of MHV infections in mice. Five-week-old female C57BL/6J mice were inoculated intranasally with either the Y, NuU, or JHM variant strain of MHV and housed with two naive mice. After 4 weeks, the presence or absence of anti-MHV antibody in the mice was determined by the enzyme-linked immunosorbent assay. We also examined the distribution of MHV in the organs of Y, NuU, or JHM variant-infected mice. Our data suggest that the transmissibility of MHV is correlated with viral growth in the gastrointestinal tract of infected mice. To the best of our knowledge, this is the first report to address the basic reproduction numbers among pathogens in laboratory animals.

## Introduction

The microbiological control of laboratory animals is essential to obtain reproducible and stable experimental results, and mouse hepatitis virus (MHV) is a representative pathogen for microbiological monitoring.^1^ MHV is a single-stranded plus-sense RNA virus with petal-like projections that belongs to the family *Coronaviridae*, and its natural host is mice.^2^ MHV causes hepatitis, enteritis, and encephalitis in mice, and its pathogenesis varies depending on the viral strain, infectious dose, route of infection, and the genetic background, age, and immune status of the host.^3-5^ Experimental infection with MHV has been used to generate hepatitis^6,7^ and a demyelinating disease mouse model..^8-11^ Studies have also been conducted to elucidate the replication and multiplication mechanisms of coronaviruses using MHV as a model.^12,13^ In recent years, the worldwide coronavirus disease 2019 pandemic caused by severe acute respiratory syndrome coronavirus 2 (SARS-CoV-2) has led to the use of MHV as a surrogate for SARS-CoV-2 in research on disinfectants.^14,15^

In Europe and the United States, MHV has the second highest infection rate in laboratory animals after mouse norovirus and mouse parvovirus.^16^ The spread of MHV in animal research facilities causes wasting disease and death in immunocompromised mice such as nude mice and suckling mice,^17,18^ and subclinical infection in adult mice reportedly modifies experimental performance in many experimental models.^19^ MHV is generally excreted in the feces of infected mice and is transmitted oronasally, but it is also transmitted via aerosols and contaminated bedding.^20^ Depending on the mouse strain and immune status, MHV-infected mice excrete infectious MHV for several days to several weeks.^21^ Contamination by MHV in a specific pathogen-free laboratory animal facility requires total culling of the infected mouse colony and disinfection of the facility, and the impact is not small.^22^ The basic principle of MHV infection control in facilities housing laboratory animals is to prevent exposure of the animals to MHV. Adequate quarantine is necessary when introducing new animals from other facilities. In addition, wild mice infected with MHV,^23^ pet mice,^24,25^ and biological materials obtained from infected mice are also sources of infection and should be handled with care.^26,27^ It is possible that the virus may be introduced into the facility unknowingly by scientists or caretakers who come into contact with MHV-contaminated animals or breeding equipment.

Serological methods such as the enzyme-linked immunosorbent assay (ELISA) and indirect fluorescent antibody methods are commonly used to diagnose MHV in animal testing facilities, and RT-PCR^28^ and RT-nested PCR methods have also been used in recent years.^29,30^

Pathogenicity and organ affinity vary among the many strains of MHV^2,31^. In this study, we used the Y, NuU, and JHM variant strains. The Y strain was isolated from suckling mice with symptoms of acute cecal colitis;^32^ the NuU strain is a less pathogenic strain that was isolated from nude mice with wasting disease;^33^ and the JHM strain was isolated from suckling mice with diarrhea.^34^ The JHM strain is used to produce a model of multiple sclerosis because it causes demyelinating encephalitis when inoculated into the brain of mice.^35^ However, the JHM strain induces acute fatal encephalitis after intracerebral infection.^3,36^ Therefore, we used the JHM variant 2.2-V-1 in this study, which was selected with monoclonal antibody J.2.2, and it loses the ability to cause acute encephalitic illness after intracerebral inoculation.^36^ When MHV is inoculated intranasally into mice, it is believed to multiply in nasal epithelial cells and spreads to other organs via the olfactory nerve, lymphatic system, and viremia.^37^

Although there have been reports comparing pathogenicity, physicochemical properties,^38^ and gene sequences of MHV viral strains,^39,40^ no direct comparison of transmissibility has been conducted. To gain a better understanding of MHV epidemiology, we attempted to determine the basic reproduction numbers of three strains of MHV in mice. The basic reproduction number (R_0_) is defined as the average number of secondary infections generated by one infected individual in a population in which all individuals are susceptible.^41,42^ We also examined viral distribution in mice infected with each MHV for a better understanding of the dynamics of MHV infection in mice. The results of this study suggest that viral growth in the gastrointestinal tract plays an important role in the transmissibility of MHV.

## Materials and Methods

### Mice

Five-week-old, MHV-free, female C57BL/6J (B6) mice were purchased from Japan SLC (Shizuoka, Japan). Some mice were inoculated intranasally with MHV and kept in cages in a negatively pressured isolator (KIS-145; Ishihara Corporation, Osaka, Japan) in the animal room, which was maintained at a temperature of 23 ± 5°C, humidity of 55 ± 5%, and 12-h illumination (light period: 8:00–20:00; dark period: 20:00–8:00). A plastic mouse breeding cage (220 × 160 × 125 mm; Ishihara Corporation) with a stainless-steel wire top was used. Each cage was filled with approximately 1,000 mL bedding (ALPHA-dri; Shepherd Specialty Papers, Watertown, TN, USA). To avoid artificial transmission of MHV, the bedding was not changed during the experiment. Mice were fed γ-ray-sterilized pellets (CRF-1; Oriental Yeast Co., Ltd., Tokyo, Japan) and tap water *ad libitum* from a plastic bottle. The animal protocol was reviewed and approved by the Institutional Animal Committee for Use and Care at the Graduate School of Agricultural and Life Sciences (University of Tokyo, Tokyo, Japan), and was conducted in accordance with the Animal Experiment Implementation Regulations and Animal Experiment Implementation Manual of the University of Tokyo.

### Viruses and cells

Y,^32^ NuU,^33^ and JHM variant (2.2-V-1)^36^ strains were cultured with DBT cells, which are MHV-sensitive.^43^ The JHM variant 2.2-V-1 was a kind gift from Dr. John O Fleming (University of Southern California School of Medicine, Los Angeles, CA then). DBT cells were cultured in Eagle’s minimal essential medium (E-MEM) containing 5% fetal bovine serum and 1% tryptose phosphate broth (Sigma-Aldrich Co., St. Louis, MO, USA) at 37°C in 5% CO_2_ in humidified air.

### Viral infection experiments

To determine the basic reproduction numbers of three stains of MHV in mice, the concentration of the Y, NuU, and JHM variant strains was adjusted to 1 × 10^4^ PFU/0.02 mL saline solution and inoculated intranasally using a micropipette into 5-week-old female B6 mice under isoflurane inhalation anesthesia. Each inoculated mouse was maintained for 2 days in a cage. Then, one inoculated mouse was bred with two naïve 5-week-old, female B6 mice for 4 weeks. Four cages were prepared for the Y strain, and five cages were prepared for the NuU and JHM variant strains. Subsequently, all mice were euthanized. Serum was collected and frozen at −80°C until use. To determine viral growth in each organ in mice, B6 mice were inoculated intranasally with 1 × 10^4^ PFU/0.02 mL of each of the Y, NuU and JHM variant strains. Four mice each were euthanized and necropsied on days 1, 3, and 5 after inoculation, and the brain, liver, jejunum, ileum, and colon were aseptically sampled and frozen at −80□ until use.

### Serological tests

The anti-MHV antibody titer in mouse serum was determined by ELISA using a commercially available ELISA kit (MONILISA® MHV 96-well; Wakamato Pharmaceutical Co., Ltd., Tokyo, Japan), and absorbance was measured using a plate reader. Transmission from MHV-inoculated mice to naïve mice was judged by seroconversion against MHV in naïve mice housed together with virus-inoculated mice.

### Quantification of infectious virus

Quantification of infectious virus in organs was performed by the plaque assay.^44^ Briefly, tissue samples from the brain, liver, jejunum, ileus, and colon were homogenized in chilled E-MEM to generate a 10% solution and then centrifuged at 3,000 rpm for 10 min. Ten-fold serial dilutions were prepared, and each dilution was assayed for infectious viruses in duplicate in DBT cells. For samples from the jejunum, ileus, and colon, 5 mg/mL gentamicin sulfate (Wako Pure Chemical Industries, Ltd., Tokyo, Japan) was added to E-MEM when generating the suspensions.

### Calculation of basic reproduction numbers

In the mouse cohabitation experiment, the average number of antibody-positive mice derived from one infected mouse was calculated to be the basic reproduction number.

### Statistical analyses

The data obtained in the experiments were subjected to statistical analyses by the Wilcoxon rank-sum test, with P < 0.05 considered statistically significant.

## Results

### Basic reproduction number of three MHV strains

To determine the basic reproduction numbers of three strains of MHV, naïve mice were cohabitated with mice inoculated with either the Y, NuU, or JHM variant strain for 4 weeks, and anti-MHV antibody production was examined by ELISA. In all four cages administered the Y strain, all naïve mice cohabitating with the mouse inoculated with the Y strain were seroconverted (Fig. 1). For the NuU strain, one naïve mouse out of two in two cages produced anti-MHV antibodies. All naïve mice in the remaining three cages were negative. Among the five cages administered the JHM variant strain, three inoculated mice showed a positive antibody response while the remaining two did not. Neither naïve mouse cohabitating with JHM variant-infected mice were seroconverted. The basic reproduction number was calculated after excluding the cages in which the inoculated mice did not show a positive antibody response. As shown in Table 1, the basic reproduction numbers of the Y, NuU, and JHM variant strains were determined to be ≥ 2, 0.4, and 0, respectively.

**Figure 1.**
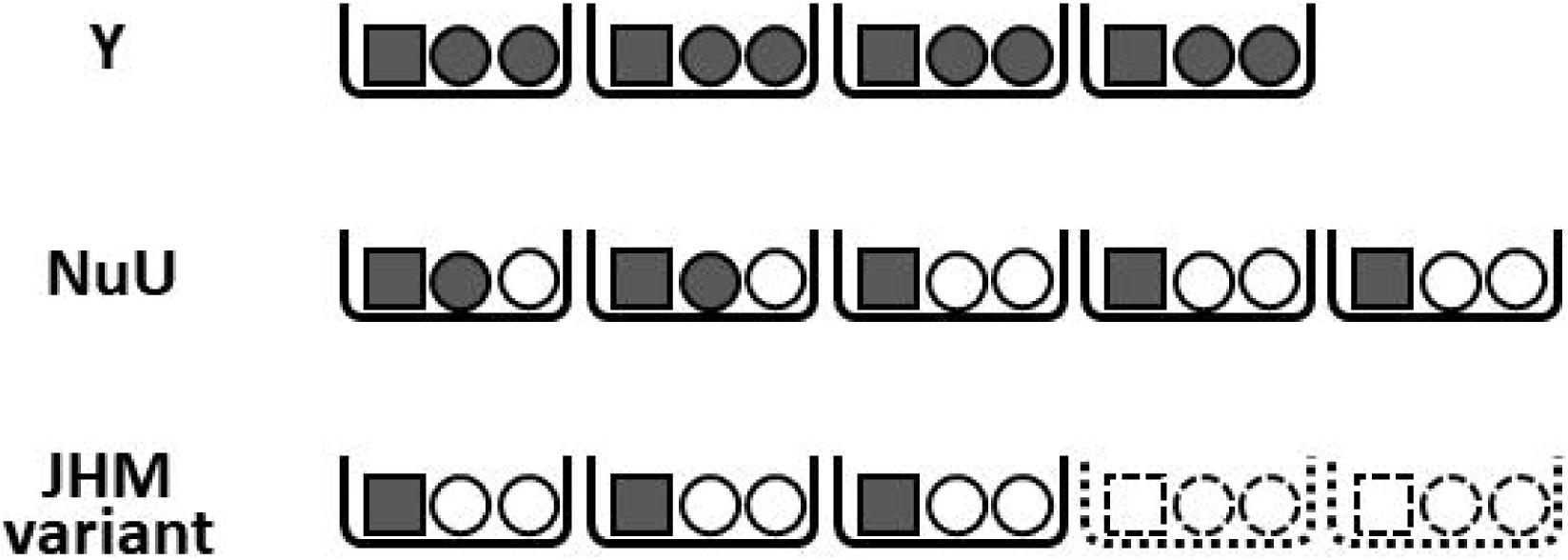
Transmission of three strains of MHV in mice by cohabitation in a cage. Four, five, and five B6 mice were inoculated intranasally with either the Y, NuU, or JHM variant of MHV, respectively. Two days later, each mouse was cohabitated with two naïve B6 mice in a cage and bred for 4 weeks. Sera were removed, and the anti-MHV antibody response was examined by ELISA. Grey and white colors indicate a positive and negative response, respectively. The square and circle indicate the inoculated and naïve mice, respectively. Since two mice inoculated with the JHM variant had a negative antibody response (shown by dotted line), data from these cages were excluded in the calculation of the basic reproduction number.

**Table 1.** Basic reproduction number of three strains of MHV

| Viral strain | Basic reproduction number |
| --- | --- |
| Y | $\geq 2$ |
| NuU | 0.4 |
| JHM variant | 0 |

### Viral growth in organs in Y, NuU, or JHM variant-infected mice

To investigate the relationship between the basic reproduction number and viral growth in various organs in mice, viral growth in the brain, liver, jejunum, ileus, and colon was examined. Five-week-old, MHV-free female B6 mice (n = 4 or 5) were inoculated with 1 × 10^4^ PFU of the Y, NuU, or JHM variant strains, and the brain, liver, jejunum, ileus and colon were removed on days 1, 3, and 5 after inoculation. In mice inoculated with the Y and NuU strains, infectious virus was detected in all organs examined (Table 2, Fig. 2). On the other hand, in mice inoculated with the JHM variant strain, infectious viruses in the jejunum, ileus, and colon were all below the detection limit. The amount of infectious virus detected in the brains of mice inoculated with the Y, NuU, and JHM variant strains tended to increase with each day of inoculation during the experiment (Fig. 2). On the other hand, there was no change in the mean amount of infectious virus detected in the liver of mice on days 3 and 5 after inoculation with the Y, NuU, and JHM variant strains. In mice inoculated with the Y and NuU strains, infectious virus was detected in the jejunum and ileus from day 1 after inoculation. In mice inoculated with the Y and NuU strains, the detection rate of infectious virus up to 3 days after inoculation was higher in the jejunum than in the ileus (Table 2). In mice inoculated with the Y and NuU strains, the detection rate of infectious virus in the colon by day 5 after inoculation was less than 50%, which was the lowest among the organs examined (Table 2).

**Table 2.**
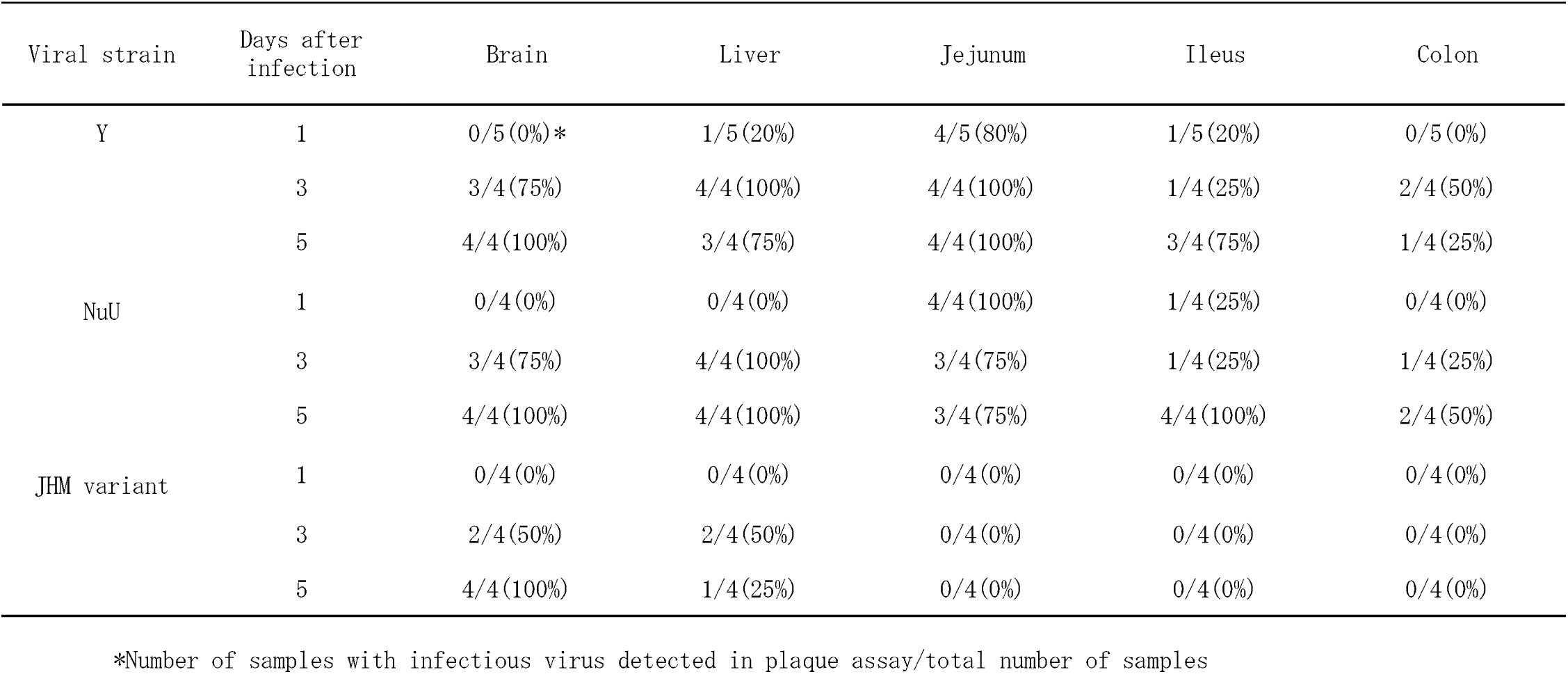
Detection rates of infectious viruses in various organs in mice inoculated intranasally with three strains of MHV

**Figure 2.**
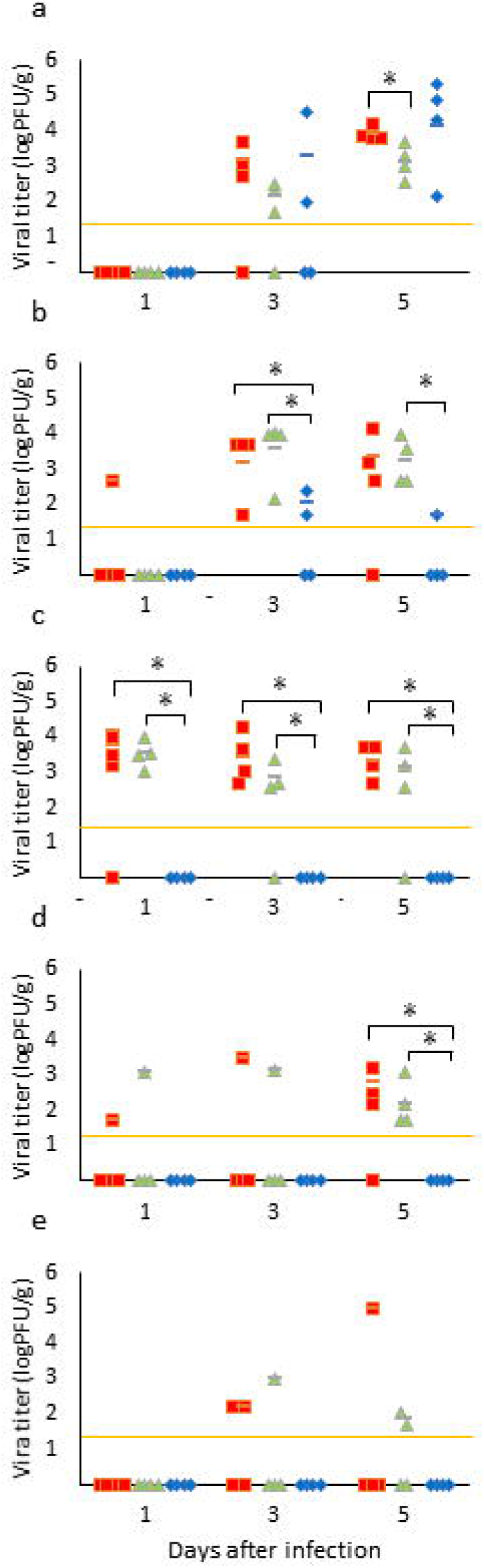
Viral growth in the organs of mice inoculated with the Y, NuU, and JHM variants. Viral titers of the brain (a), liver (b), jejunum (c), ileus (d), and colon (e) in mice inoculated intranasally with the Y 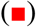, NuU 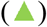, or JHM variant (⍰) of MHV was measured on days 1, 3, and 5 after infection. The horizontal black bar indicates the average of four samples. The detection limit of MHV in the plaque assay is indicated by the orange line. □: P < 0.05

## Discussion

Basic reproduction number is an important epidemiological parameter defined as the average number of secondary infections generated by one infected individual in a population in which all individuals are susceptible.^41,42^ Basic reproduction number is not simple and is affected by numerous biological, sociobehavioral, and environmental factors that govern pathogen transmission.^45^ In this study, we attempted to determine the basic reproduction number of MHV in mice in laboratory animal facilities. The data suggest that the Y strain is the most transmissible, and the JHM variant is not transmissible by cohabitation. There have been some reports addressing how MHV spreads among mice in laboratory animal facilities,^46,47^ but to the best of our knowledge, no study has examined the basic reproduction number of MHV in mice.

Infectious virus was detected in all organs of mice inoculated with Y and NuU strains, while infectious virus in the jejunum, ileus, and colon of mice inoculated with JHM variant was below the detection limit. The difference in transmissibility between the Y and JHM variants may be because the JHM variant is less likely to increase in the gastrointestinal tract and be excreted in the feces compared with the Y strain in the case of intranasal inoculation. The quantitative results of infectious virus in organs did not explain the difference in transmissibility between the Y and NuU strains. For both strains, examination of the amount of virus excreted in feces and excretion period using another assay, for example, quantitative PCR is expected to provide evidence of a difference in transmissibility. In addition, considering the infection route of MHV, we could not exclude the possibility of physicochemical stability of each viral strain in the external environment where mouse feces are discharged.

Barthold *et al*^48^. detected infectious virus in the gastrointestinal tract at 3 and 5 days after intranasal inoculation of BALB/cByJ mice with 10^3^ of the median tissue culture infectious dose of the JHM strain. However, in this study, no infectious virus was detected in the gastrointestinal tract of C57BL/6J mice intranasally inoculated with the JHM strain. There are many substrains of JHM with different antigenicities.^49^ The JHM strain used in this study is a variant strain resistant to S protein-specific monoclonal antibody^36^ and is not identical to the strain used by Barthold *et al*.^48^ The reason for the different results from those of Barthold *et al*. may be due to the difference in viral strain and host (BALB/cByJ vs. B6).

The cage and rack system have been developed and marketed for the purpose of better microbiological control of laboratory animals. MHV is a representative pathogen for mice and is used for the evaluation of system functions.^50,51^ Based on our results, it may be possible to evaluate the protective function of cages and racks against MHV infection in mice by using a strain with relatively high transmissibility, such as the Y strain.

## Acknowledgements

We thank Dr. John O. Fleming for providing a JHM variant (2.2-V-1).

